# The molecular basis of parental conflict driven regulation of endosperm cellularization

**DOI:** 10.1101/2023.06.22.546051

**Authors:** N. Butel, Y. Qiu, W. Xu, J. Santos-González, C. Köhler

## Abstract

The endosperm is a seed tissue supporting embryo growth, similar to the placenta in mammals. It originates after fertilization of the maternal central cell by one of the paternal sperm cells. In the early stages of *Arabidopsis thaliana* seed development, nuclei divisions in the endosperm are not followed by cellularization. After a defined number of mitotic cycles, the endosperm cellularizes and stops dividing. The timing of endosperm cellularization impacts on final seed size and is differentially controlled by maternal and paternal genome contributions. While increased maternal genome dosage causes early endosperm cellularization and the formation of small seeds, the opposite is caused by increased paternal genome dosage. The parental factors controlling the differential timing of endosperm cellularization remain largely unexplored. Here, we show that a family of maternally expressed auxin response factors (*ARFs*) promotes endosperm cellularization and regulates final seed size.

**One-Sentence Summary:** Endosperm cellularization is under antagonistic parental control that converges on Auxin Response Factors.

## Introduction

The endosperm is a reproductive tissue derived from the fusion of a haploid sperm cell with a predominantly diploid central cell, which sustains and supports embryo development^1^.

In *Arabidopsis thaliana*, like in most angiosperms, endosperm development occurs in two phases. In the initial phase, endosperm nuclei proliferation is not followed by cellularization, resulting in the formation of a coenocyte^2^. At a tightly controlled timepoint, a wave of cellularization starts from the micropylar region surrounding the embryo to reach the opposite chalazal endosperm^2^. At the end of the process, most of the endosperm is cellularized and nuclear divisions cease. The timing of the transition from the first to the second phase is critical for seed development. Precocious or delayed cellularization leads to very small or enlarged seeds of impaired viability, respectively^3^. Endosperm cellularization is under differential parental control; while increased maternal genome dosage promotes cellularization, increased paternal genome dosage has the opposite effect by delaying cellularization.

Previous work identified auxin as a critical factor initiating the first nuclear divisions of the endosperm and determining the timing of endosperm cellularization^4,5^. Auxin biosynthesis is initiated after fertilization from the paternal genome by *YUCCA10* (*YUC10*) and *TRYPTOPHAN AMINOTRANSFERASE RELATED 1* (*TAR1*), two imprinted paternally expressed genes regulating auxin biosynthesis^4^. Auxin levels cease at the time of cellularization, while conversely, endosperm cellularization failure correlates with increased auxin levels^4^. How auxin controls endosperm cellularization is nevertheless unknown.

## Results

We identified a cluster of Auxin Response Factors (ARFs) that is strongly upregulated in seeds with delayed endosperm cellularization^4,5^. Given the connection between auxin and endosperm cellularization, we investigated the function of those ARFs in the endosperm.

This ARF cluster contains eight members that are located in the pericentromeric region of chromosome 1 (Fig. 1A). All members share high sequence similarity, indicating that they function redundantly (Fig. S1). The exception is *ARF23*, which is truncated and has been proposed to be a pseudogene and was therefore not considered further^6^. We will refer to these centromeric *ARFs* as *cARFs*.

**Fig. 1.**
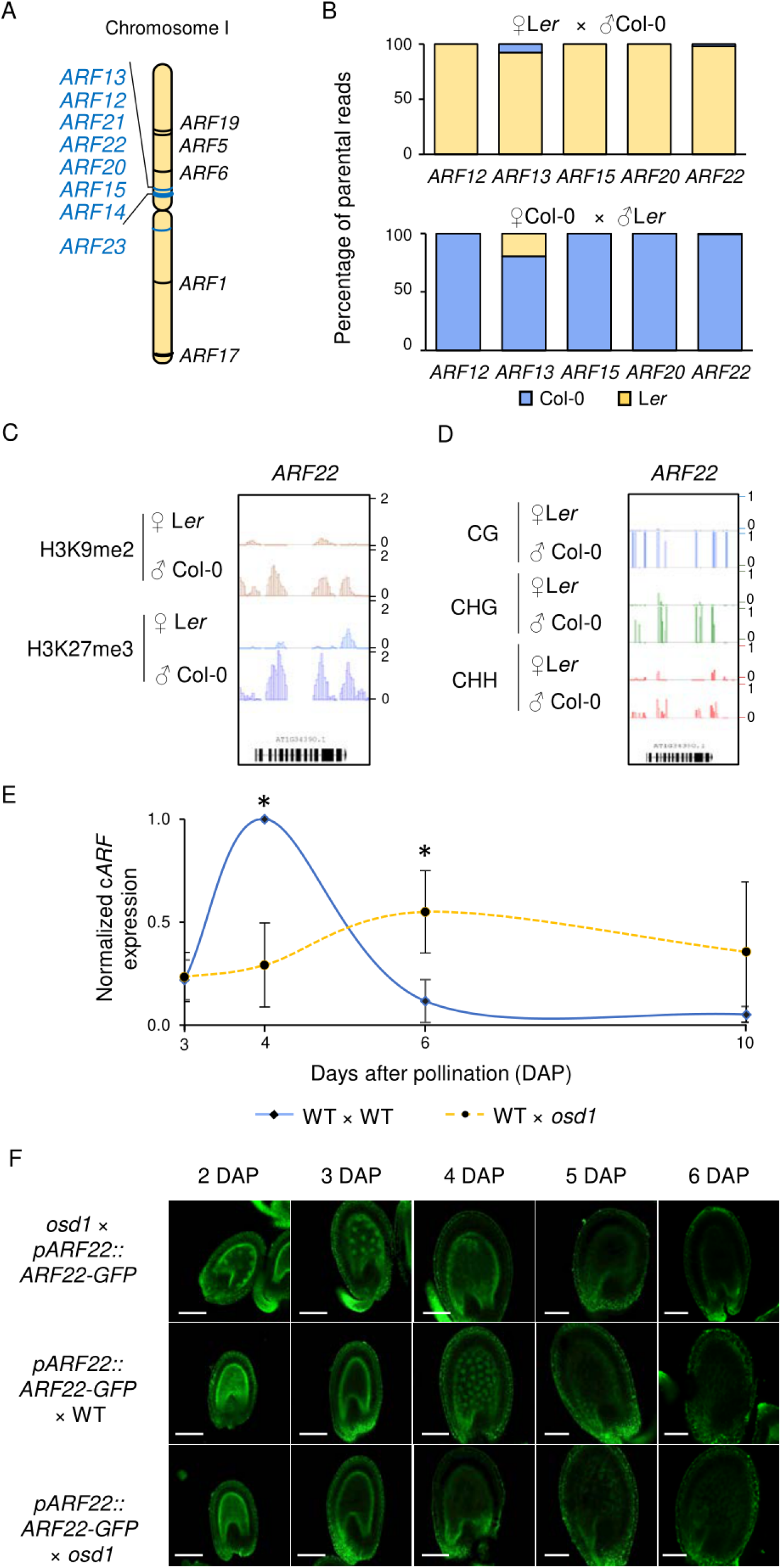
*cARFs* are expressed at the onset of endosperm cellularization. (**A**) Localization of Arabidopsis *ARF* genes on chromosome 1. *cARFs* are indicated in blue. (**B**) Percentage of parental *cARF* reads derived from crosses of Col-0 and L*er* accessions in the 4 DAP endosperm^7^. (**C**) Parental-specific enrichment of H3K9me2 (red) and H3K27me3 (blue) histone marks on *ARF22* in the 4 DAP endosperm^22^. (**D**) Parental-specific DNA methylation on *ARF22* in the endosperm at 6 DAP^23^ (**E**) qRT-PCR analysis of *cARF* expression in 3-, 4-, 6- and 10 DAP siliques of the indicated crosses. Errors bars represent the standard deviation of five independent biological replicates. Asterisks mark statistically significant differences based on Student’s t-test (p<0.05) (**F**) Confocal microscopy pictures showing expression of *pARF22:ARF22- GFP* at different stages of seed development in the indicated crosses. Scale bars, 100μm.

Based on available transcriptome data of the endosperm 4 days after pollination (DAP)^7^, *ARF12, ARF13, ARF15, ARF20* and *ARF22* are maternally expressed genes (MEGs), thus the maternal alleles are exclusively or preferentially expressed in the endosperm (Fig. 1B). *ARF14* and *ARF21* are also expressed in the endosperm at 4 DAP, but the imprinting status could not be determined due to either missing sequence polymorphisms (SNPs) between Col-0 and L*er* accessions in the case of *ARF21* or reads covering SNPs in the case of *ARF14*. The paternal alleles of all *cARFs* are highly DNA methylated and enriched for repressive histone methylation on H3 lysine 9 and lysine 27 (H3K27me3 and H3K9me2, respectively), correlating with the specific silencing of the paternal alleles (Fig. 1C and D, and Fig. S2A and B).

### cARFs are expressed at the onset of endosperm cellularization

To specifically determine when and where *cARFs* were expressed, we monitored transcript abundance by qRT-PCR and protein localization using a reporter construct for ARF22, which contained the promoter and coding region of *ARF22* fused to the green fluorescent protein (*GFP*) reporter (*pARF22::ARF22-GFP*) (Fig. 1E and F). *cARF* transcript levels peaked at 4 DAP and similarly, GFP fluorescence accumulated in both the micropylar and the peripheral endosperm at around 4-5 DAP (Fig. 1E and F). Thus, cARF accumulation preceded endosperm cellularization, which in Arabidopsis wild-type Col-0 initiated at 5-6 DAP.

In seeds inheriting a double dosage of paternal chromosomes (referred to as paternal excess crosses), *cARFs* were deregulated^5^, suggesting that *cARFs* are sensitive to parental genome dosage. To test this hypothesis, we monitored *pARF22::ARF22-GFP* expression in seeds with unbalanced parental genome dosage. We made use of the *omission of second division 1 (osd1)* mutant that produces 2n male and female gametes at high frequency^8^. Thus, using *osd1* as either female or male parent, allowed generating seeds with either increased maternal or paternal genome dosage, correlating with precocious (4-5 DAP) or delayed endosperm cellularization (after 6 DAP), respectively^3^.

We found that increased paternal genome dosage generated by crossing wild-type (WT) plants with *osd1* pollen donors caused reduced and delayed *cARF* transcript accumulation, shifting the peak of expression from 4 DAP to 6 DAP (Fig. 1E). This pattern was also reflected by the *pARF22::ARF22-GFP* reporter; we did not detect GFP fluorescence in paternal excess seeds between 2 DAP and 6 DAP (Fig. 1F).

Conversely, in maternal excess seeds where *osd1* was the female parent, *ARF22-GFP* expression could be already detected at 2-3 DAP (Fig.1F). This early expression was not a consequence of increased copy number, since the *pARF22::ARF22-GFP* reporter was not imprinted and introduced through pollen. We failed to detect *ARF22* transcripts by qRT-PCR, likely because the number of endosperm nuclei was too few to allow detecting low abundant endosperm transcripts at this developmental stage. Nonetheless, the detection of precautious *ARF22-GFP* activity strongly suggests that *cARF* expression is sensitive to the maternal genome dosage and that increased maternal genome dosage correlates with increased *cARF* expression.

Together, these results show that *cARF* expression is antagonistically regulated by maternal and paternal genome dosage, reflecting their MEG identity. Furthermore, *cARF* activity correlates with the onset of endosperm cellularization^3^, suggesting a functional role of cARFs in regulating this process.

### cARF deficiency delays endosperm cellularization

Single T-DNA insertions in *ARF15, ARF20* and *ARF22* did not cause abnormalities in seed development, suggesting functional redundancy of cARFs (Fig. S3). Using Crispr/Cas9 with two guide RNAs targeting multiple *cARFs*, we identified one line with premature stop codons in *ARF13* and *ARF20* (Fig. 2A).

**Fig. 2.**
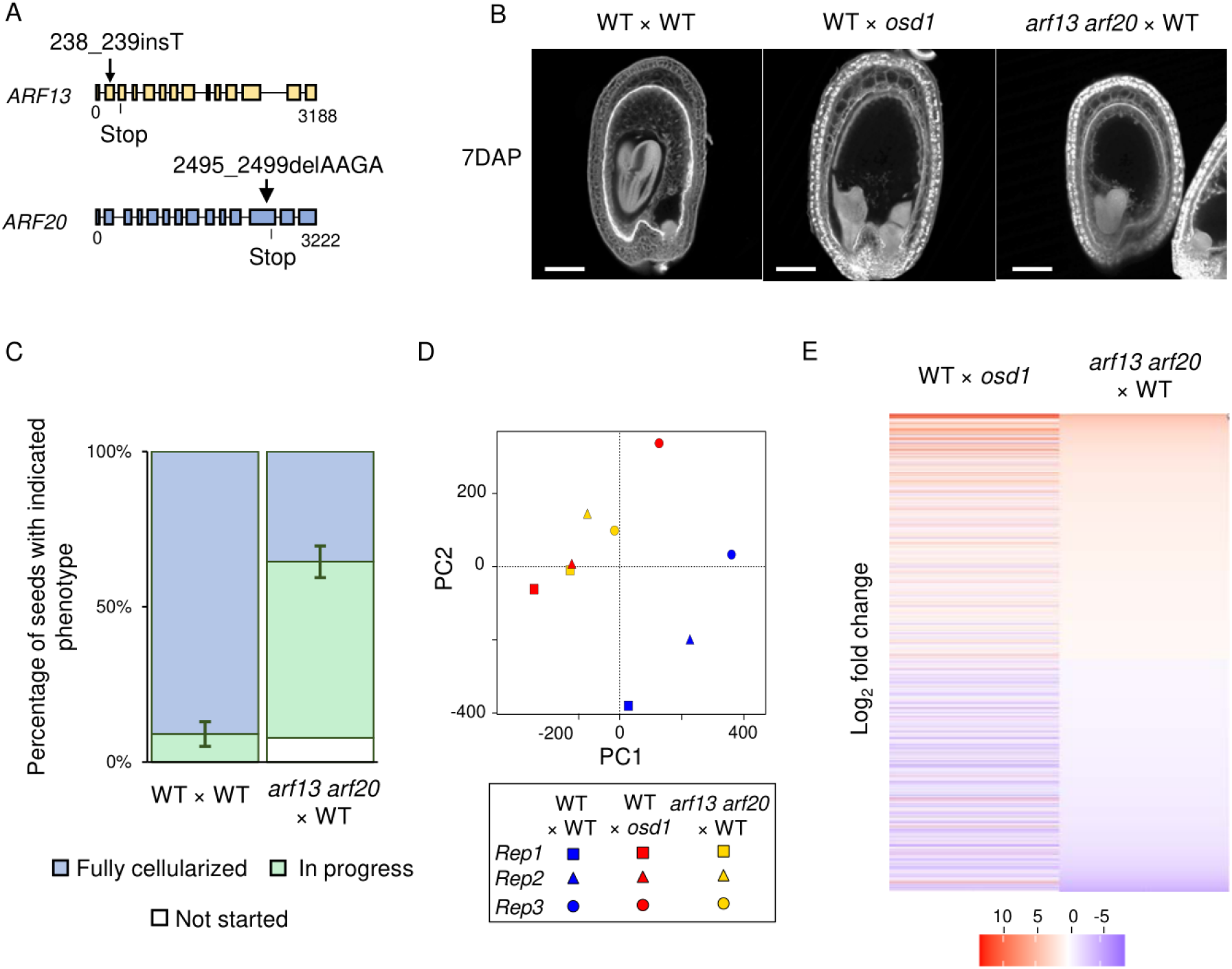
Mutations in *cARFs* delay endosperm cellularization. (**A**) Schematic representation of *ARF13* and *ARF20* and positions of two mutations induced by CRISPR/Cas9. The filled squares correspond to exons. (**B**) Multiphoton microscopy pictures of 7 DAP Feulgen stained seeds derived from indicated crosses. Scale bars, 100μm. (**C**) Quantification of endosperm cellularization in seeds of indicated crosses. In progress refers to seeds where cellularization has initiated but not terminated. Error bars represent the standard deviation of two independent biological replicates where at least 50 seeds were analyzed per replicate. (**D**) Principal Component Analysis of transcriptomes of 7 DAP seeds of the indicated genotypes. (**E**) Heatmap showing the log2 fold change of deregulated genes in *arf13 arf20* × WT compared to WT and WT × *osd1* compared to WT at 7 DAP. Only those genes are shown that were significantly deregulated in WT x *osd1* compared to WT (|Log2FC| >= 1 ; padj<0.05).

Since *ARF13* and *ARF20* are predominantly maternally expressed, we pollinated *arf13 arf20* with WT pollen to test the effect on endosperm cellularization. Loss of maternal ARF13 ARF20 function delayed endosperm cellularization; while all wild-type seeds were completely cellularized at 7DAP, the majority of *arf13/+ arf20/+* seeds were uncellularized or had only started the cellularization process, resembling paternal excess seeds (Fig. 2B and C). These results reveal that maternal *cARFs* have a functional role in endosperm cellularization and likely induce cellularization.

To test whether the delay of endosperm cellularization in *arf13 arf20* and paternal excess seeds has a common molecular basis, we compared the transcriptomes of seeds lacking maternal ARF13 ARF20 function with paternal excess seeds at 7DAP, when the corresponding wild type was fully cellularized. We found indeed that the transcriptomes of paternal excess seeds and *arf13 arf20* seeds clustered together, whereas the wild-type transcriptomes clustered separately (Fig.2D). The similarity in transcriptomes was also reflected by a similar trend of deregulated genes in paternal excess seeds and seeds lacking ARF13 ARF20 function (|log2Fc| >= 1; padj < 0.05) (Fig.2E).

Together, the transcriptional response in seeds lacking ARF13 and ARF20 function resembled that of paternal excess seeds, supporting the hypothesis that delayed cellularization in paternal excess seeds is linked to the misregulation of *cARFs*.

### cARF overexpression induces early cellularization

We next addressed the question whether precocious expression of *cARFs* is sufficient to induce early cellularization and thus mimic a maternal excess seed phenotype. To this end we expressed *ARF22* in the endosperm under control of the *PHERES1 (PHE1)* promoter that is active directly after fertilization and lasts until completion of endosperm cellularization (Fig. S4).

Consistent with the idea that cARFs are required to induce endosperm cellularization, *pPHE1::ARF22* lines produced seeds with precociously cellularized endosperm, preceding wild-type seeds by one or even two days (Fig. 3A and Fig. S5A). Hemizygous *pPHE1::ARF22* lines produced aborted seeds at high frequency (40% to 60%, Fig. 3B and C) revealing that precocious expression of *ARF22* is sufficient to trigger seed arrest. Those seeds contained well developed embryos surrounded by a small, cellularized endosperm, similar to maternal excess seeds^9^ (Fig. 3A, and Fig. S5A and B). Together, this data show that induction of endosperm cellularization correlates with *ARF22* expression.

**Fig. 3.**
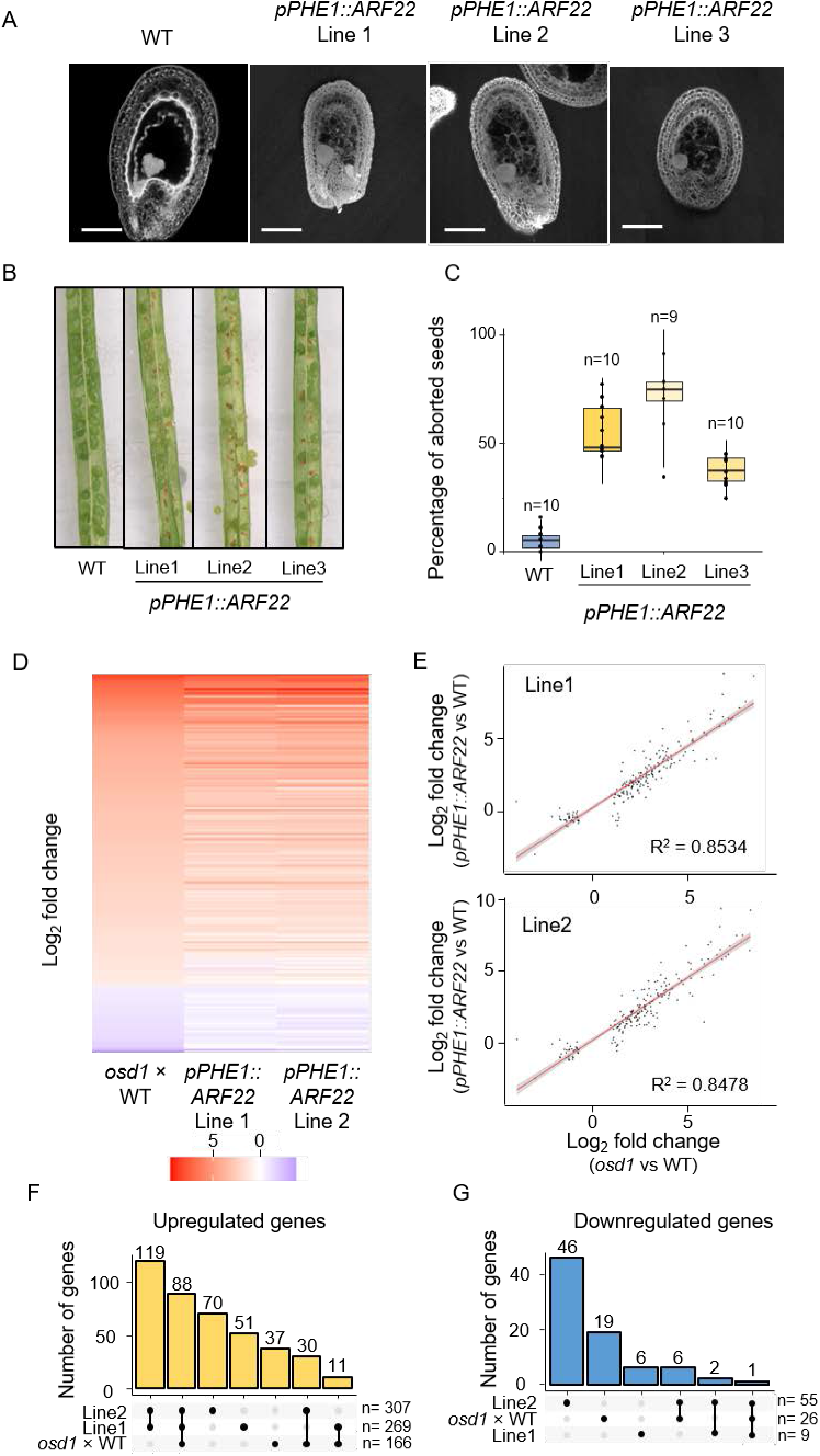
Precocious *cARF* expression promotes endosperm cellularization. (**A**) Multiphoton microscopy pictures of 5 DAP Feulgen stained seeds of three independent *pPHE1::ARF22* lines. Scale bars, 100μm. (**B**) Pictures showing seed abortion in the *pPHE1::ARF22* lines. (**C**) Quantification of seed abortion in three independent *pPHE1::ARF22* lines. Each dot corresponds to the percentage of aborted seeds in one silique. The number of analyzed siliques is indicated on the top. (**D**) Heatmap showing the log2 fold change of deregulated genes in 4 DAP seeds of *osd1* × WT compared to WT and *pPHE1::ARF22* compared to WT. Only genes are shown that were significantly deregulated in *osd1* × WT compared to WT (|Log2FC| >= 1 ; padj<0.05). (**E**) Correlation plot of log2 fold changes of deregulated genes in *pPHE1::ARF22* lines and the *osd1* × WT crosses. The linear regression is shown in red and the coefficient of correlation R^2^ is indicated in the chart. (**F, G**) Upset plot showing the number of commonly upregulated (F) and downregulated (G) genes in the different transcriptomes.

Strikingly, expression of *pPHE1::ARF22* did not only change the time of endosperm cellularization, but also affected the pattern of this process. In wild-type seeds, endosperm cellularization started at the micropylar region surrounding the embryo and spread from there over the whole endosperm (Fig. S6). In contrast, in *pPHE1::ARF22* lines, cellularization started at both ends simultaneously and the generally uncellularized chalazal endosperm became completely cellularized (Fig. S5A and Fig. S7). This cellularization pattern corresponds with the activity of the *PHE1* promoter, which is strongly expressed in the chalazal region of the endosperm^10^. Together, this data strongly supports the hypothesis that ARF22 directly induces endosperm cellularization.

Similar phenotypes were observed when overexpressing *ARF15* and *ARF21* under control of the *PHE1* promoter, in line with the proposed redundant function of cARFs in promoting endosperm cellularization (Fig. S7).

To test whether the phenotypic similarities between seeds overexpressing *cARFs* and maternal excess seeds was reflected at the molecular level, we compared the transcriptomes of two *pPHE1::ARF22* lines with maternal excess seeds (*osd1* × WT) at 4 DAP. At this timepoint cellularization had not yet started in WT, but was completed in the other genotypes (Fig. 3A). Significantly deregulated genes (|log2Fc| >= 1 ; padj < 0.05) in maternal excess seeds were similarly deregulated in seeds of *pPHE1::ARF22* lines, corresponding to a strong correlation between the datasets (Fig. 3D and E). The majority (78%) of upregulated genes in maternal excess seeds were also upregulated in at least one of the *pPHE1::ARF22* lines and about half (53%) of them were commonly upregulated in both lines (Fig. 3F and G). The 88 commonly upregulated genes were enriched for functions related to phragmoplast and cytoskeleton fiber formation, consistent with the induced cellularization process (pvalue<0.05).

Together, our data uncover cARFs as key regulators of endosperm cellularization that act in a dosage dependent manner and likely underpin the parental dosage sensitivity of endosperm cellularization.

#### Evolution of *cARFs* in angiosperms

Phylogenetic analysis revealed that Arabidopsis *cARFs* are derived from a Brassicaceae-specific duplication of *ARF9* (Fig. S8A and B) and in many Brassicaceae crown species the ancestral *cARFs* duplicated into tandem arrays nested in pericentromeric regions (Fig. S8C). The recurring copy number increase of *cARFs* and the conserved location in pericentromeric heterochromatin suggests selection towards increased maternal-specific expression of *cARFs* in the Brassicaceae.

The *cARFs* are more similar to the tandem paralogs within a species than to orthologs in sister species (Fig. S8C), suggesting that frequent events of gene conversion homogenized the cluster of *cARFs*^11,12^. Concerted evolution of *cARFs* leading to multiple copies of nearly identical *cARF* genes may have evolved as a mechanism allowing maternal control of endosperm cellularization. This evolutionary pattern is consistent with the predictions of the parental-conflict theory^13,14^, which forecasts the evolution of maternally expressed suppressors of endosperm growth to counteract paternally expressed growth promoters^15^.

The *ARF9* clade arose from the γ-whole genome triplication shared by all core eudicots^14^, while the paralogous clade corresponds to *ARF11/18*^14^ (Fig. S8B). The identified ortholog of *ARF9/11/18* in maize, *ZmARF7* (Zm00001eb118970), is expressed in the endosperm sharply around the cellularization stage, putatively promoting the transition from the nuclear to the cellular phase^16^. We thus speculate that the repressive *ARF* clade harboring the *cARFs* and *ARF9/11/18* play a conserved role in promoting endosperm cellularization. In line with this hypothesis, the orthologs of *ARF9/11/18* in several species are also expressed in the early endosperm or seed transcriptomes (Fig. S8B). In contrast, Arabidopsis *ARF9/11/18* are not expressed in the early endosperm (Fig. S4), suggesting that the rise of *cARFs* allowed them to adopt specialized functions in the endosperm. The loss of a broad expression pattern may have promoted the increase of copy number without detrimental effects on sporophyte development.

## Discussion

The timing of endosperm cellularization is decisive for final seed size and a major target of parental conflict^17^. Our study reveals that parental-dosage dependent regulation of *cARFs* controls endosperm cellularization, implicating cARFs as molecular targets of parental conflict (Fig. 2, 3 and 5).

cARFs belong to the evolutionary conserved ARF B class that are considered to be transcriptional repressors^18,19^. Repressive B class ARFs were shown to antagonize activating A class ARFs^20^, providing an intuitive model whereby cARFs block auxin-mediated endosperm proliferation^4^ by competing with activating A-type ARFs that remain to be identified (Fig. 4A).

**Fig. 4.**
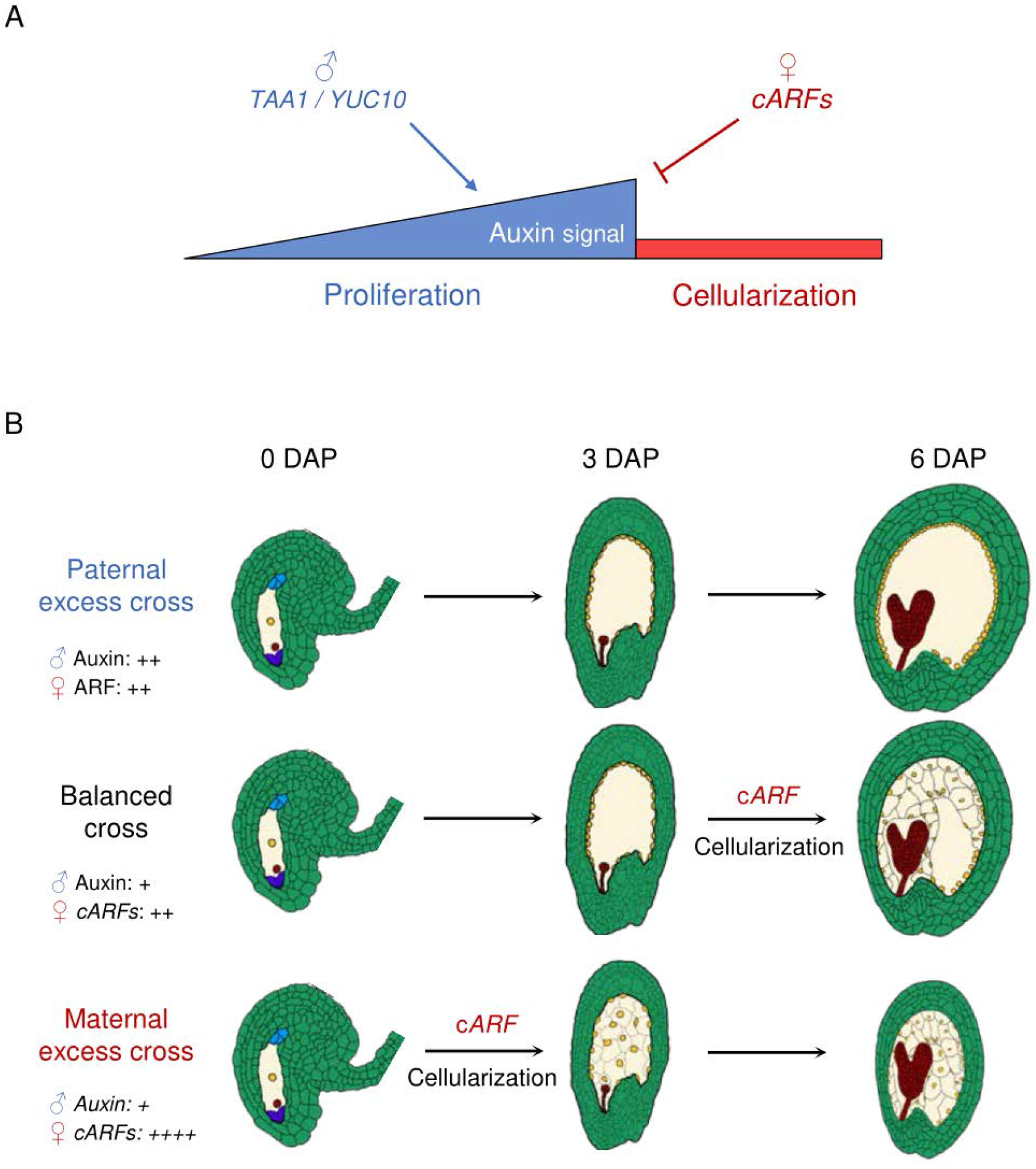
Model depicting antagonistic parental effects on endosperm cellularization via regulation of auxin production and signalling. **(A)** After fertilization, the paternally expressed genes *YUC10* and *TAA1* trigger auxin production and initiate endosperm proliferation. Proliferation ends when *cARFs* are expressed from the maternal genome and likely block auxin signaling, thereby inducing endosperm cellularization. **(B)** Altering the parental genome dosage changes the time of *cARF* accumulation and endosperm cellularization. In paternal excess crosses, the double dosage of the paternal genome stimulates auxin production, reducing the effect of maternally produced *cARF* transcripts, leading to a delay or absence of endosperm cellularization. Conversely, in maternal excess crosses, doubling of the maternal genome causes increased accumulation of *cARF* transcripts, precociously reaching the threshold to induce cellularization.

A key prediction of the parental conflict theory is that maternal and paternal genomes antagonistically affect the growth of embryo supportive tissues^15^. Specifically, natural selection is expected to favour paternally-active alleles promoting seed growth and maternally-active alleles restricting seed growth. By promoting endosperm cellularization and thus restricting seed growth, cARFs are likely major targets of this conflict. Consistent with the predictions regarding maternally-biased expression of growth suppressors, *cARFs* are maternally expressed while paternally silenced by a combination of repressive epigenetic modifications (Fig. 1C and D, and Fig. S2). Interestingly, within the Brassicaceae we found evidence for a repeated amplification of *cARFs* into tandem arrays nested in pericentromeric regions (Fig. S8B). This recurring copy number increase of *cARFs* is likely a consequence of parental conflict, ensuring maternal control of endosperm cellularization. Auxin biosynthesis in the endosperm is controlled by the paternal genome and increased auxin levels delay endosperm cellularization^5^, revealing an antagonistic parental control of endosperm cellularization converging on auxin biosynthesis and signaling (Fig. 4B).

In conclusion, we identified cARFs as maternally active dosage-sensitive regulators of endosperm cellularization. cARFs induce endosperm cellularization and thus restrict seed growth, making them direct molecular targets of parental conflict in angiosperm seeds.

## Acknowledgments

We thank the Green Team and the Microscopy facility of the Max Planck Institute of Molecular Plant Physiology for supporting this work.

## Funding

Knut and Alice Wallenberg Foundation grant 2018-0206 (CK)

Knut and Alice Wallenberg Foundation grant 2019-0062 (CK)

Max Planck Society, Germany.

## Author contributions

Conceptualization: NB and CK

Methodology: NB and CK

Investigation: NB(Experiments), YQ (phylogenetic analysis), WX (CRISPR/Cas9 design)

Software: JSG

Visualization: NB, YQ, JSG

Funding acquisition: CK

Project administration: CK

Supervision: NB and CK

Writing – original draft: NB, YQ and CK

Writing – review & editing: NB, YQ, WX and CK

## Data and materials availability

RNA-seq data generated in this study is available at NCBI’s Gene Expression Omnibus database, under the accession number GSE232803. The imprinting, CHiP-seq, DNA methylation and endosperm expression data can be found under the GSE119915^21^, GSE66585^22^, GSE84122^23^ and GSE12404^24^ respectively.

## Supplementary Materials

Materials and Methods

Figs. S1 to S8

Tables S1

Data S1 to S3

References

