## Supplemental material and figures for "The molecular basis of parental conflict driven regulation of endosperm cellularization"

**The PDF file includes:**

Materials and Methods
Figs. S1 to S8
Tables S1
References

**Other Supplementary Materials for this manuscript include the following:**

Data S1 to S3

### Material and Methods

#### *Plant cultivation and lines used in this study*

The Arabidopsis mutant *osd1-3* has been previously characterized[1]. The *arf15-1* (SALK_029838) and *arf20-2* (SALK_032522) mutants have been published[2]. The *arf22-3* (SALKseq_49790) mutant has been characterized in this study. Primers used to genotype the mutants are listed in Supplemental table 1. For all experiments, Col-0 was used as the wild-type control.

Arabidopsis seeds were sterilized for 15min in a solution of 70% ethanol and 0,0001% Triton X-100 and washed with 100% ethanol for an additional 15 min. Dried seeds were sown on plates containing ½ Murashige and Skoog medium and stratified at 4°C for two days. Plates were incubated in a growth chamber for two weeks (16 h light/8 h dark; 60µmol/s/m2; 22°C), then transferred to soil and grown in phytotron chambers (16 h light/8 h dark; 150µmol/s/$m^2$; 21°C; 70% humidity).

#### *Generation of plasmids and transgenic plants*

Genes were amplified from Arabidopsis Col-0 genomic DNA with the primers described in Supplemental Table 1. After amplification, the fragments were inserted into a pENTR vector by using the pENTR/D-TOPO kit (ThemoFisher Ref: K240020SP). For the *pPHE1::cARFs,* the fragments were inserted into the *pPHE1-pB7WG2*[3] vector using a LR reaction (ThermoFisher Ref: 11791020). For the *pARF22::ARF22-GFP* construct, the destination vector was *pB7FWG.0*.

For the CRISPR construct, the guideRNA sequences for mutating *cARFs* were designed by E-CRISP[4]. Two guide RNAs were chosen to target cARF genomic DNA: DT1(AAGTTTATTACTTTCCTCAAGGG) and DT2c (AAAGATCCCATTGAAGAAATTGG). The construction protocol has been previously published[5,6]. The PCR fragment was amplified from pCBC-DT1T2 with the four primers listed in (Supplemental Table 1) and inserted into pHEE401E by Golden Gate cloning.

All constructs were introduced into the Arabidopsis Col-0 accession using the floral dip protocol[7]. Transformed plants were selected on medium containing appropriate chemicals.

#### *Microscopy*

For monitoring *pARF22::ARF22-GFP*, siliques were opened at the indicated stage and seeds were mounted in water. Fluorescence was observed using a LEICA Stellaris 8 Dive microscope with an emission of 488 nm and an excitation range of 493-551 nm.

For clearing and Feulgen staining, siliques were opened at indicated stages and incubated overnight at 4°C in a fixing solution of ethanol: acetic acid (3:1). On the next day, the solution was replaced by 70% ethanol and stored at -20°C until staining.

For seed clearing, the seeds were removed from the siliques and incubated overnight at 4°C in a clearing solution (66.7% [w/w] chloralhydrate, 8.3% [w/w] glycerol). They were then mounted in clearing solution and observed on an Olympus BX-51 microscope.

Sample preparation and embedding for Feulgen staining was done as previously described [3]. Samples were observed on a LEICA Stellaris 8 Dive microscope using the multiphoton mode with an emission of 400 nm and with an excitation of 563-668 nm.

#### *RNA extraction, qRT-PCR and library preparation*

For qRT-PCR, two siliques were harvested at the indicated stage, ground in liquid nitrogen and stored at -80°C until extraction. For mRNA sequencing, around 500 seeds were dissected from siliques and stored in RNA*later* solution (ThermoFisher Ref: AM7021) at 4°C before extraction.
RNA was extracted using the RNeasy Plant mini Kit (Qiagen Ref: 74904). RNAs were treated by DNAseI at 37°C for 30 min (ThermoFisher: EN0521). DNAseI was inactivated by an incubation at 65°C for 10min and, removed by a TRIzol extraction prior to library construction following the manufacturer's protocol (ThermoFisher Scientific, catalog no. 15596018).
The reverse transcription reaction was performed using the RevertAid H Minus First Strand cDNA Synthesis Kit (ThermoFisher Ref: K1631) and a dTTTN primer (see Supplemental Table 1). The qPCR was performed with the Power SYBR™ Green PCR Master Mix (ThermoFisher Ref: 4367659) and the indicated primers (Supplemental Table 1). Normalization of cARF transcripts was done relative to GAPDH.
The mRNA libraries were generated using the NEBNext® Ultra™ II DNA Library Prep Kit (NEB Ref: E7645S) coupled to the NEBNext® Poly(A) mRNA Magnetic Isolation Module (NEB Ref: E7490S). Sequencing was done by Novogene on a HiSeqX in 150-bp paired-end mode.

**RNAseq analysis**
For each replicate, 150 bp long paired-end reads were trimmed using Trimgalore (5 bp at the 5' end and 20 bp at the 3' end) and mapped to the Arabidopsis (TAIR10) genome using hisat2. Mapped reads were counted using Htseq-count and normalized to Transcripts Per Million (TPM) for genes using StringTie. Differentially regulated genes between conditions and across the replicates were detected using DESeq2 applying a threshold of log2 (foldchange)>=1 with a false discovery rate (FDR) adjusted P-value of <0.05. A multivariate analysis (Principal Component Analysis) was performed in order to assess the replicability and degree of similarity between samples using the vegan package in R.

**Phylogenetic analyses**
To elucidate the relatedness within the ARF family, amino acid sequences of all 23 ARFs in Arabidopsis were obtained from TAIR10. MUSCLE was used to generate the multiple sequence alignments with default settings[8]. The sequences of the three defining functional domains, B3 type DNA-binding domain (InterPro entry: IPR003340), auxin response factor domain (IPR010525) and AUX/IAA domain (IPR033389), were identified by the conserved domain search tool, CD-Search[9], and were extracted and aligned independently, to generate the concatenated alignments of conserved ARF protein regions. IQ-TREE 1.6.7 was applied for maximum likelihood inference of phylogeny[10], with the JTT substitution model as suggested by the implemented ModelFinder[11] and one thousand ultrafast bootstrap replicates to estimate the support for reconstructed branches[12].
To analyze the phylogenetic timing of *cARF* and *ARF9* duplication, amino acid sequences of homologs of *ARF9*, *ARF11* and *ARF18* were identified in several angiosperm species, with an emphasis on Brassicales (Supplemental Data S3). Full-length sequence

alignments by MUSCLE were used as input for the IQ-TREE analyses, following the procedure above.

To investigate the pattern of cARF evolution after the divergence from ARF9, amino acid sequences and nucleotide sequences of cARFs and ARF9 in several Brassicaceae species (Supplemental Data S3) were used to generate a guided codon alignment by MUSCLE. A maximum likelihood tree was then generated by IQ-TREE with the codon alignment as the input, the GTR substitution model and one thousand replicates of ultrafast bootstrap.

A

Coding regions

| | *ARF12* | *ARF13* | *ARF14* | *ARF15* | *ARF20* | *ARF21* | *ARF22* | *ARF23** |
|---|---|---|---|---|---|---|---|---|
| *ARF12* | 100% | 50% | 87% | 90% | 87% | 90% | 92% | 88% |
| *ARF13* | | 100% | 51% | 51% | 51% | 50% | 51% | 56% |
| *ARF14* | | | 100% | 87% | 84% | 86% | 88% | 87% |
| *ARF15* | | | | 100% | 90% | 92% | 91% | 88% |
| *ARF20* | | | | | 100% | 92% | 89% | 87% |
| *ARF21* | | | | | | 100% | 91% | 85% |
| *ARF22* | | | | | | | 100% | 86% |
| *ARF23** | | | | | | | | 100% |

**Fig. S1. cARFs share high sequence similarity.**

Percentage of identity at the protein level between each cARF. *ARF23* is a pseudogene, indicated by the asterisk.

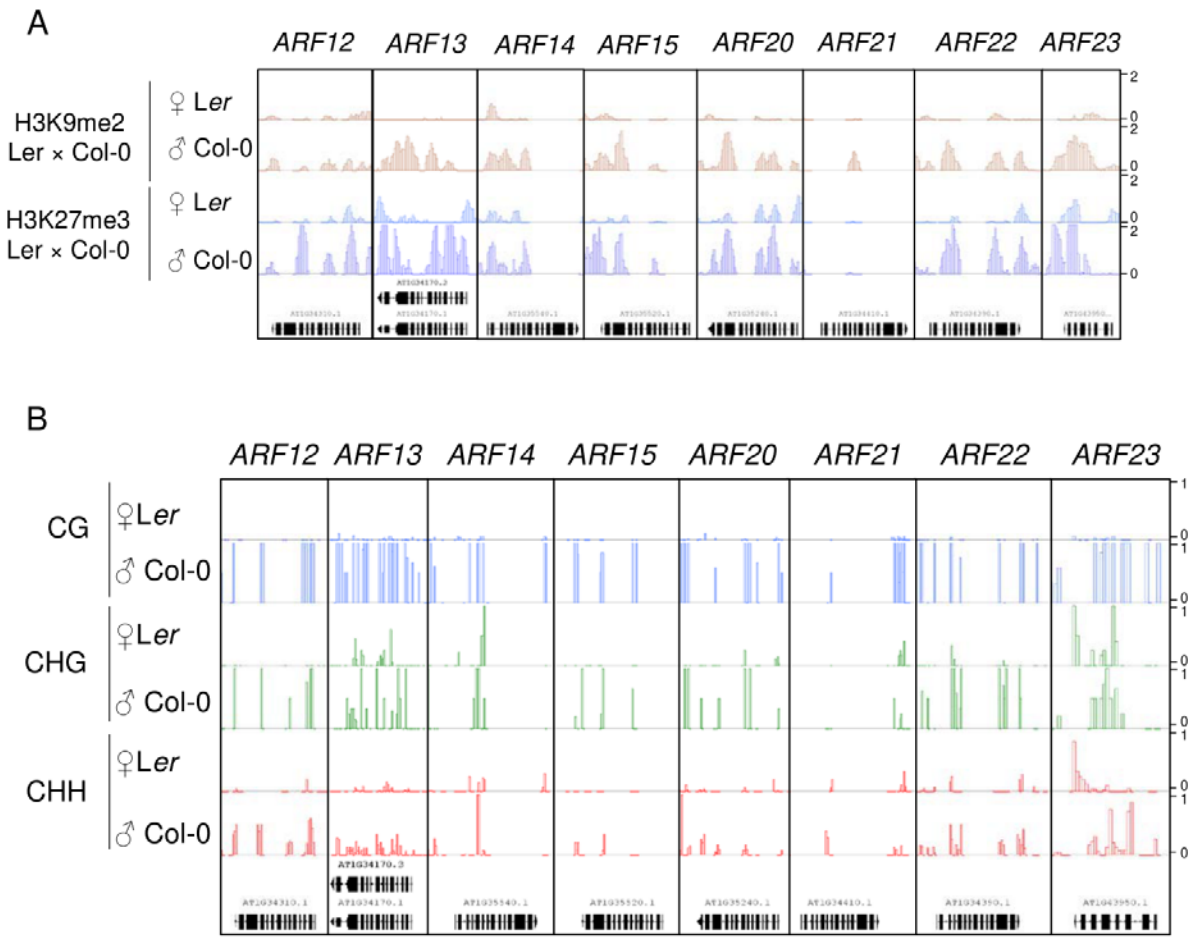


**Fig. S2. The paternal allele of *cARFs* is marked by repressive histone modifications and DNA methylation.**

(**A**) Parental-specific enrichment of H3K9me2 (red) and H3K27me3 (blue) histone modifications on *cARFs* in the 4 DAP endosperm[13]. (**B**) Bedgraphs showing parental-specific DNA methylation in the endosperm at 6DAP[14].

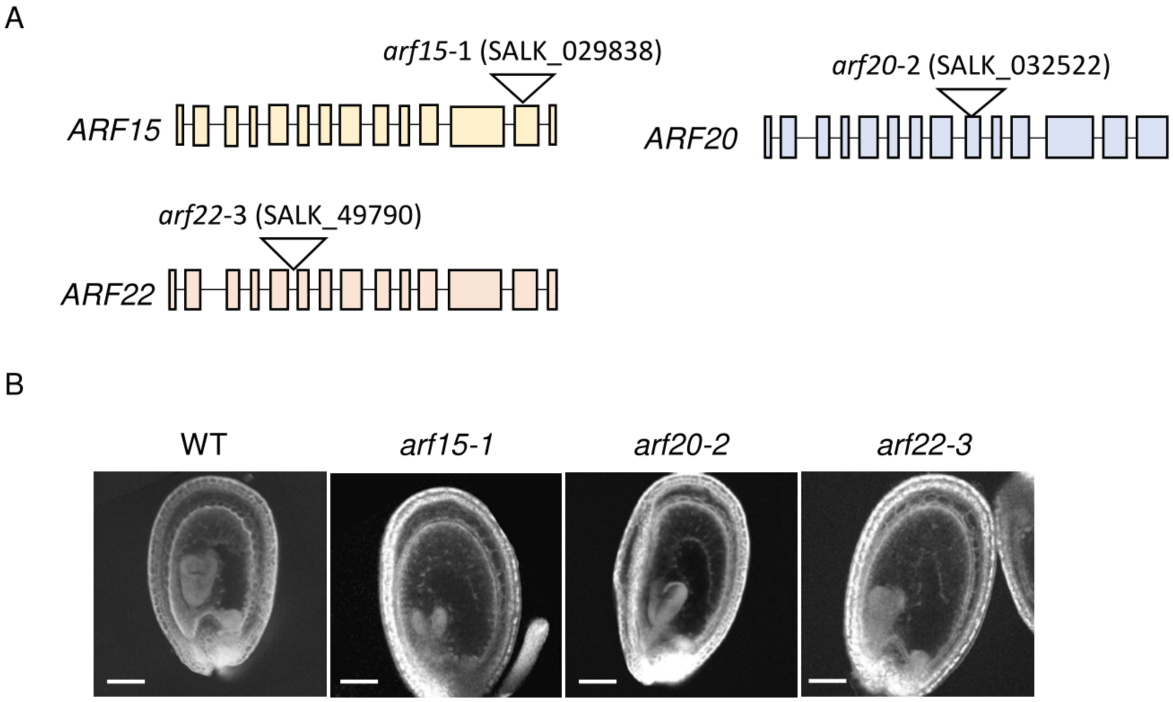


**Fig. S3. Single *arf* T-DNA insertion mutants do not exhibit abnormal seed phenotypes.**

(**A**) Schematic representation showing the position of T-DNA insertions in *ARF15*, *ARF20* and *ARF22*. Filled boxes correspond to exons. (**B**) Multiphoton microscopy pictures of 6 DAP Feulgen stained seeds. Scale bars, 100µm.

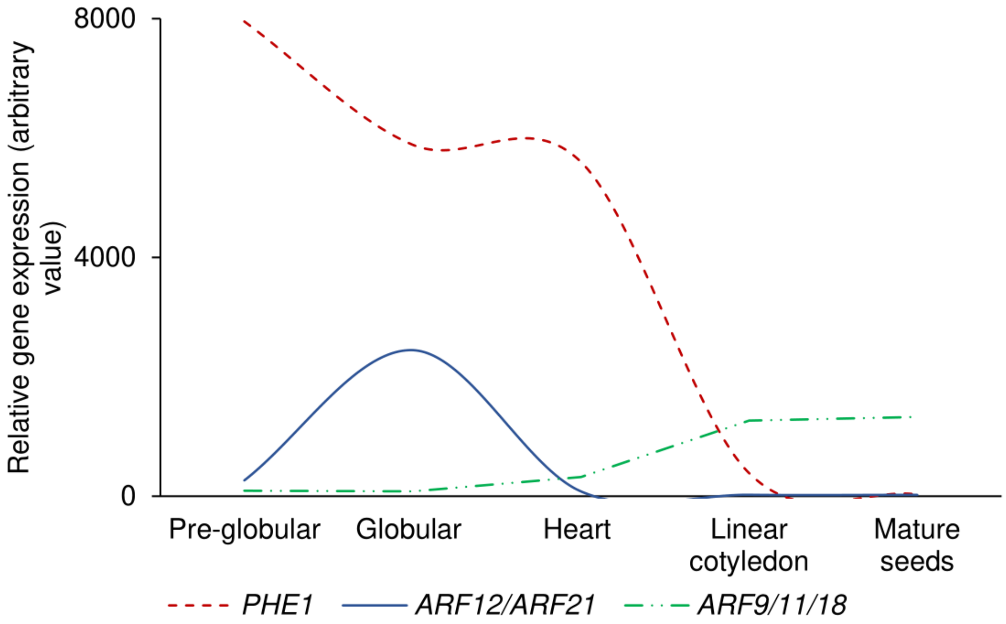


**Fig. S4. Comparison of relative *PHE1* and *cARF* expression during different stages of endosperm development.**

Expression of *PHE1, ARF9/11/18* and *cARFs* based on[15]. The data include only two of the eight *cARFs*, allowing only a comparison of the time of *cARF* expression relative to *PHE1*, but no absolute quantitative comparison.

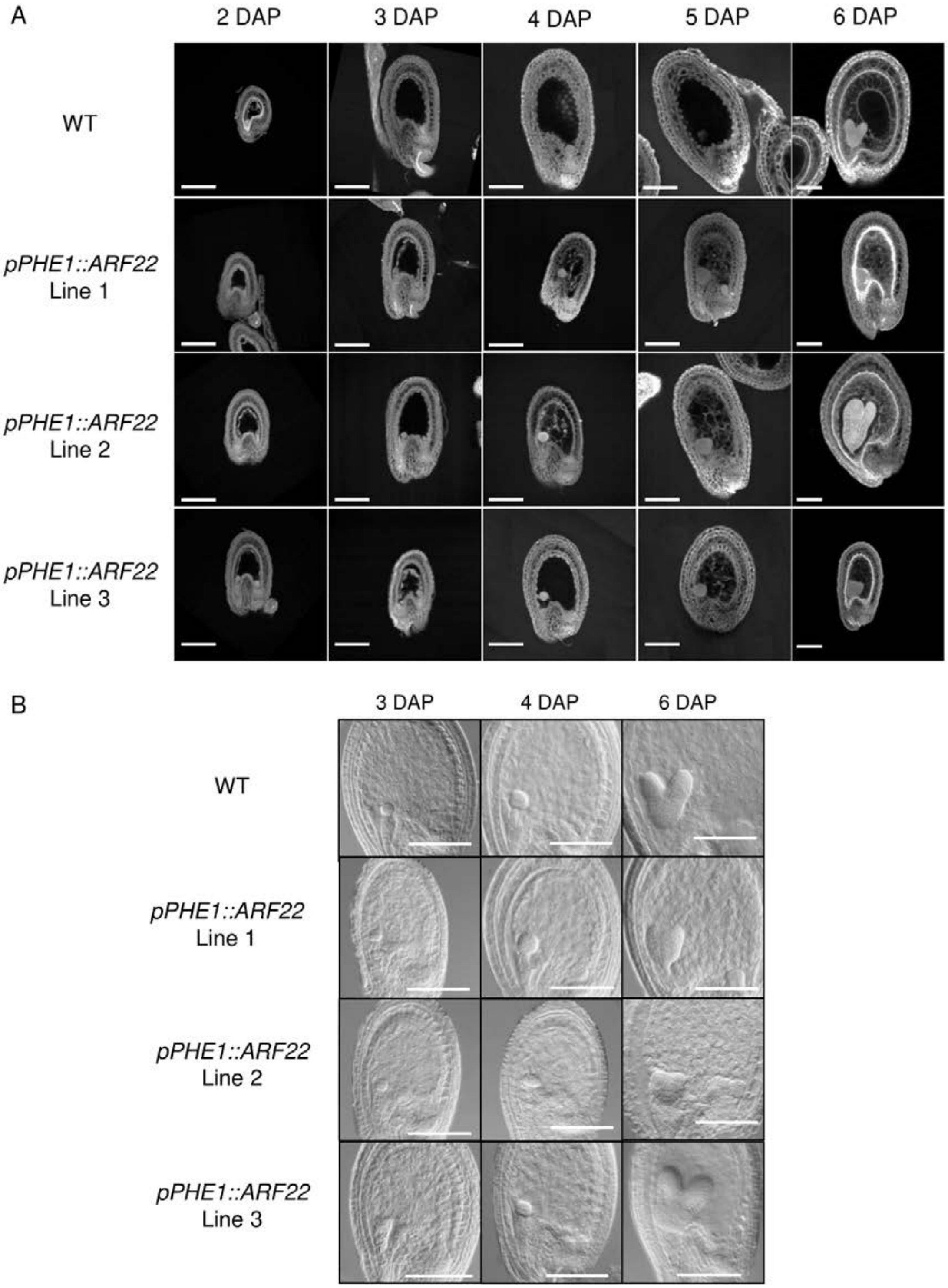


**Fig. S5. Seed phenotypes of the *pPHE1::ARF22* lines.**

(**A**) Multiphoton microscopy pictures of Feulgen stained seeds taken at the indicated time points. Scale bars, 100µm. (**B**) Pictures of cleared seeds taken at the indicated time points. Scale bars, 100µm.

A

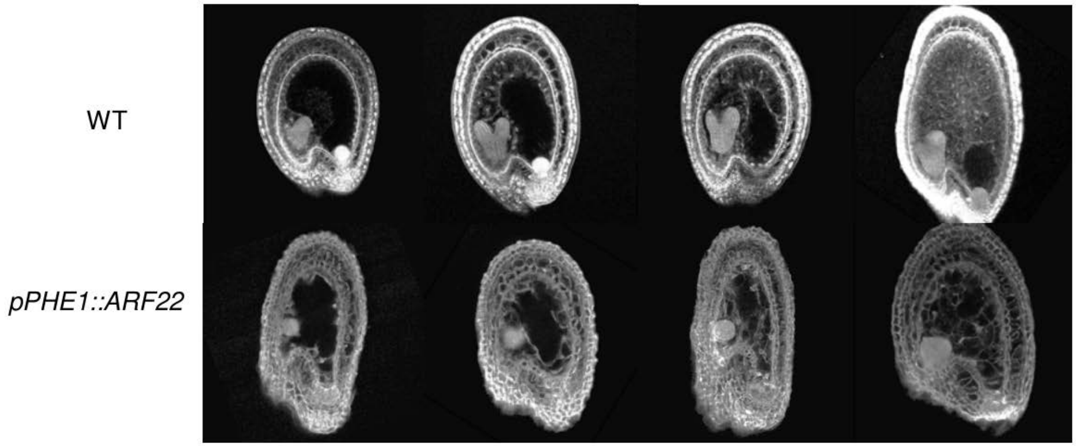


B

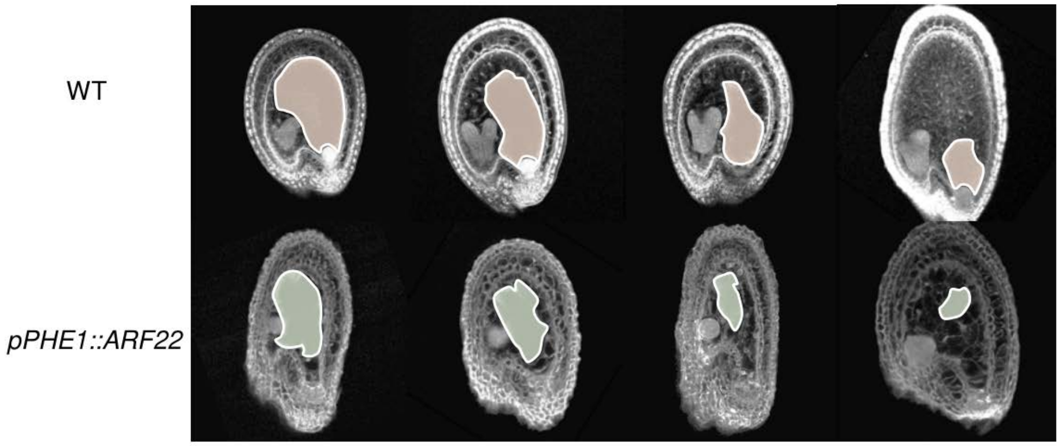


**Fig. S6. Endosperm cellularization in WT and *pPHE1::ARF22* seeds.**
(**A**) Multiphoton microscopy pictures of Feulgen stained seeds taken either at 6 DAP (WT) or at 4 DAP (*pPHE1::ARF22* seeds). (**B**) Same pictures as in (A) but with the non-cellularized endosperm indicated in brown and green for WT and *pPHE1::ARF22*, respectively.

A

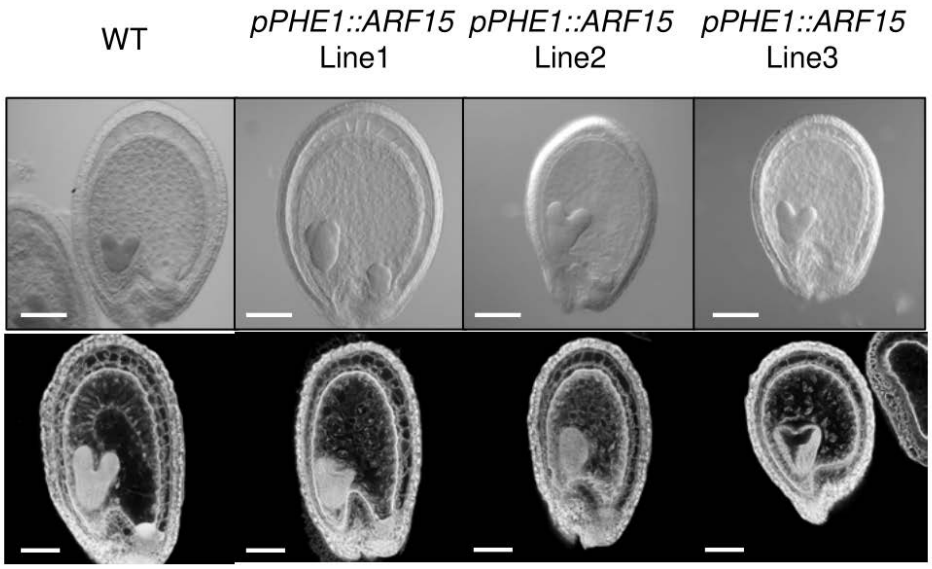


B

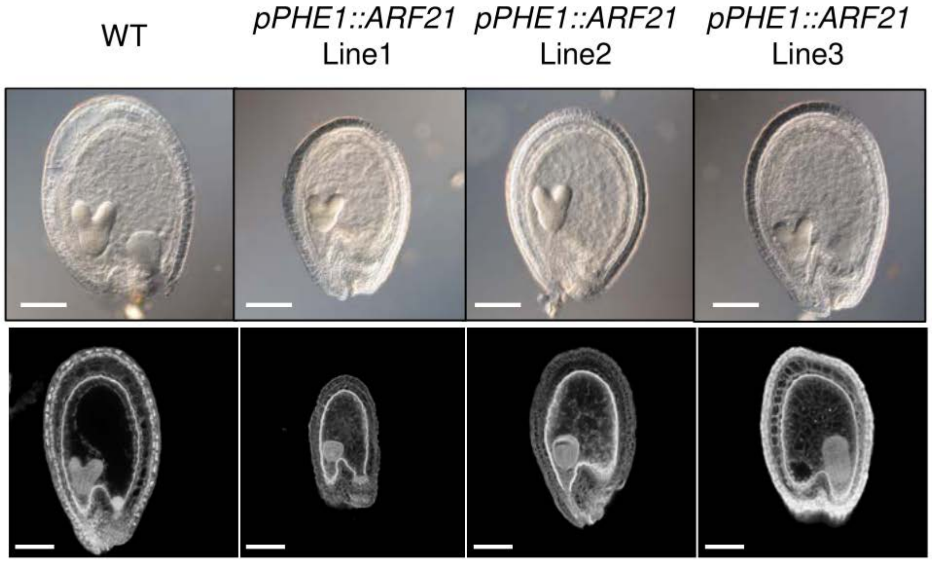


**Fig. S7. *pPHE1::ARF15* and *pPHE1::ARF21* exhibit an early endosperm cellularization phenotype.**

(**A**) DIC pictures of cleared seeds (upper part) or multiphoton microscopy pictures of Feulgen stained seeds (bottom part), of different *pPHE1::cARF* lines at 6 DAP. Three independent lines were analyzed for each construct. Scale bars, 100µm

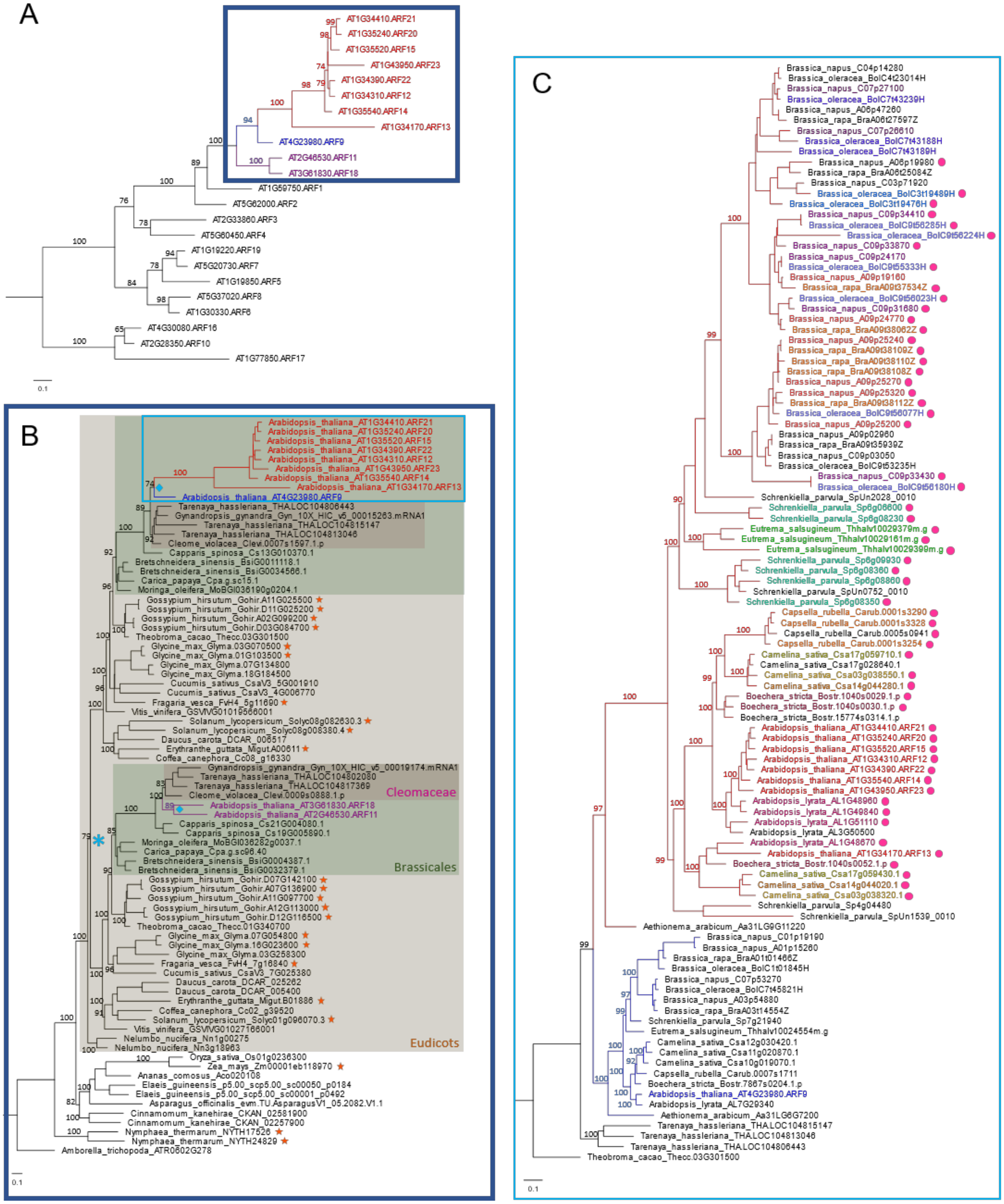


**Fig. S8. Maximum-likelihood (ML) trees of ARFs with bootstrap values supporting branches of interest.**

(A). The phylogeny of 23 ARFs in Arabidopsis showing the clade of cARFs (red), ARF9 (blue) and ARF11/18 (purple). (B). The phylogeny of cARFs (red), ARF9 (blue) and

ARF11/18 (purple) in angiosperms. The asterisk marks the eudicot γ- Whole Genome Triplication, and the diamonds mark the Brassicaceae-specific gene duplication. Genes labelled by orange stars are the ARF9/11/18 homologs with confirmed expression in early-stage endosperm or seed transcriptomes. (C). The phylogeny of cARFs (red) and ARF9 (blue) in the Brassicaceae. cARFs are colored by tandem clusters. Pink dots label ARFs located in pericentromeric regions. The source of sequences, transcriptomes and centromere locations are listed in Supplementary Data S3.

| Function | Name | Sequence | Reference |
|---|---|---|---|
| *osd1-3* genotyping | osd1-3-F | tcagttttggtctggcatgggt | - |
| | osd1-3-R | CTTCTGGATCTCCGCCATCACAAT | - |
| | osd1-3-T-DNA | CTGGGAATGGCGAAATCAAGGCATC | - |
| *arf15* genotyping | arf15-F1 | CAACTAAGCGTGGCCTCCCCGATG | - |
| | arf15-R1 | accggtaacattttcgacccc | - |
| *arf20* genotyping | arf20-F1 | CTTTGATGCCAGATACAACTgtaag | - |
| | arf20-R1 | GGCATTGGTTGGACAGGGTAG | - |
| *arf22* genotyping | arf22-F1 | acgagcttATGGAAAGTGGCA | - |
| | arf22-R1 | CCGCATTGTAAATCTTGAACC | - |
| pPHE1:ARF15 cloning | gARF15-F1 | **CACC**ccgatctttggataagaggttATGG | - |
| | gARF15-R1 | accggtaacattttcgacccc | - |
| pPHE1:ARF21 cloning | gARF21-F1 | **CACC**acgagcttATGGAAAGTGGCA | - |
| | gARF21-R1 | caccggtaacatttttgacccc | - |
| pPHE1:ARF22 cloning | gARF22-F1 | **CACC**acgagcttATGGAAAGTGGCA | - |
| | gARF22-R1 | tgagagactcTTACTGGACTTCA | - |
| pARF22:ARF22-GFP cloning | pARF22gARF22-F1 | **CACC**cgtcgtcaacctctgcgtat | - |
| | pARF22gARF22-R1 | T**A**ACTGGACTTCAAGTTTTTGACC | |
| Reverse transcriptase primer | dTTTN | TTTTTTTTTTTTTTTTTTVN | - |
| cARFs qPCR | cARF-Rt-F1 | GGCATTGGTTGGACAGGGTAG | - |
| | cARF-Rt-R1 | CAACTAAGCGTGGCCTCCCCGATG | - |
| GAPDH qPCR | GAPDH-F | GGTACGACAACGAATGGGGT | Elvirat-Matelot et al., 2016 |
| | GAPDH-R | TGACTGCGCATGGAATCAGT | Elvirat-Matelot et al., 2016 |
| CRISPR arf guide 1 RNA | ARF-DT1-BsF | ATATATGGTCTCGATTGAGTTTATTACTTTCCTCAAGTT | |
| | ARF-DT1-F0 | TGAGTTTATTACTTTCCTCAAGTTTTAGAGCTAGAAATAGC | |
| CRISPR arf guide 2 RNA | ARF-DT2c-R0 | AACATTTCTTCAATGGGATCTTCAATCTCTTAGTCGACTCTAC | - |
| | ARF-DT2c-BsR | ATTATTGGTCTCGAAACATTTCTTCAATGGGATCTTCAA | - |
| pB7WG2 modification with pPHE1 | pPHE1-F1 | GAGCTCgactttaaaatagtagaaaagcttg | Figueiredo et al., 2015 |
| | pPHE1-R1 | ACTAGTatctcttatctttttcttttgtgt | Figueiredo et al., 2015 |

### Table S1. Primers used in this study

**Data S1. Analysis of the *arf13 arf20* transcriptomes.**

(A) Table showing normalized reads for the 7 DAP seed transcriptomes. (B) and (C) Tables showing DESeq2 results comparing WT transcriptomes to Col-0 × *osd1* (B) or to *arf13 arf20* × Col-0 (C).

**Data S2. Analysis of the *pPHE1::ARF22* transcriptomes.**

(A) Table showing the normalized reads for the 4 DAP seed transcriptomes. (B), (C) and (D) Tables showing DESeq2 results comparing WT libraries to *osd1* ×Col-0 (B), to *pPHE1::ARF22* Line1 (C) or to *pPHE1::ARF22 Line2* (D).

**Data S3: Species surveyed for the phylogenetic analyses.**
